# Structural basis for constitutive activation and CXCL1 recognition of human herpesvirus 8-encoded G protein-coupled receptor KSHV-GPCR

**DOI:** 10.1101/2023.12.27.573477

**Authors:** Aijun Liu, Yezhou Liu, Clàudia Llinàs del Torrent Masachs, Weijia Zhang, Leonardo Pardo, Richard D. Ye

## Abstract

Kaposi’s sarcoma-associated herpesvirus (KSHV) encodes a viral G protein-coupled receptor, KSHV-GPCR, that contributes to KSHV immune evasion and pathogenesis of Kaposi’s Sarcoma. KSHV-GPCR shares a high similarity with CXC chemokine receptors CXCR2 and can be activated by selected chemokine ligands. KSHV-GPCR is also unique for its constitutive activity by coupling to various G proteins. We investigated the structural basis of ligand-dependent as well as constitutive activity of KSHV-GPCR through cryo-EM structural determination of KSHV-GPCR-Gi signaling complexes with and without bound CXCL1 chemokine ligand. Analysis of the apo-KSHV-GPCR-Gi structure, with an overall resolution of 2.81 Å, unraveled the involvement of extracellular loop 2 in constitutive activation of the receptor. This and other structural motifs serve to stabilize the constitutively-active KSHV-GPCR. The CXCL1-bound KSHV-GPCR-Gi structure was solved to an overall resolution of 3.01 Å, and showed a two-site binding of the chemokine by the receptor. Together with functional validations, this work shed light on the structural basis for constitutive as well as CXCL1-induced activation of KSHV-GPCR. The work also demonstrates evolutionary advantage in immune evasion by KSHV through its virally encoded chemokine receptor, with potential implications in developing therapeutic strategies for KSHV infection.

## Introduction

Kaposi’s sarcoma-associated herpesvirus (KSHV), also termed human herpesvirus-8 (HHV8), is a γ2-herpesvirus and the etiologic agent of Kaposi’s sarcoma (KS), a multifocal spindle-cell tumor (Karamanou et al., 2013; Mesri et al., 2010). In addition to KS, the virus causes primary effusion lymphoma, multicentric Castleman’s disease and KSHV inflammatory cytokine syndrome (Goncalves et al., 2017). As a double-stranded DNA virus, KSHV has a genome of around 165,000 base pairs that encode many homologs of mammalian proteins, likely pirated by the virus from its mammalian host cells (Moore et al., 1996). These hijacked genes code for functional orthologs of interleukin-6, Bcl-2, cyclin-D, chemokines CCL2 and CCL3, interferon regulatory factors and a G protein-coupled receptor (GPCR) in KSHV (De Groof et al., 2021; Rosenkilde et al., 2022). Altogether, these genes hijacked from host cells are believed to contribute to KSHV viral immune evasion.

Among the virus-encoded proteins of KSHV, KSHV-GPCR (encoded by *ORF74*) contributes to immune responses, tumor transformation and angiogenesis (Arvanitakis et al., 1997; Bais et al., 1998). KSHV-GPCR is highly expressed by the host cell during the lytic cycle (Mesri *et al*., 2010). The upregulation of KSHV-GPCR and other lytic proteins including viral interferon regulatory proteins that inhibit type I interferons, and the expression of host genes, particularly cytokines interleukin 6 and CXCL1, are upregulated by co-expression of KSHV-GPCR with cell-type specificity (Polson et al., 2002). Moreover, the expression of KSHV-GPCR also elicits VEGF-mediated angiogenesis in a paracrine manner (Cesarman et al., 2000; Masood et al., 1997). It has been suggested that the oncogenesis mechanism of KSHV highlights KSHV-GPCR, which can activate NF-κB and AP-1 signaling pathways and induce the expression of growth factors, cytokines and chemokines (Schwarz and Murphy, 2001; Shepard et al., 2001). *In vivo* studies also suggest that transgenic expressing KSHV-GPCR could cause Kaposi’s sarcoma-like tumors (Guo et al., 2003). These findings point out the ability of KSHV to modulate host responses for immune evasion (Areste and Blackbourn, 2009; Coscoy, 2007).

KSHV-GPCR shares a high sequence similarity with human CXC chemokine receptors, especially CXCR1 and CXCR2 binding with common chemokine ligands including CXCL1, CXCL2 and CXCL8 (Rosenkilde *et al*., 2022). However, KSHV-GPCR is unique for its ligand-independent constitutive activity (Arvanitakis *et al*., 1997). As a result, KSHV-GPCR may influence cellular functions similar to other chemokine receptors but without binding endogenous ligands. Several other viral GPCRs also share this striking feature of constitutive activity, including Epstein-Barr virus (EBV)-encoded BILF1, human cytomegalovirus (HCMV)-encoded US28, US27, UL33, UL78, human herpesvirus 6 and human herpesvirus 7 (HHV6/HHV7)-encoded U12 and U51 (De Groof *et al*., 2021; Rosenkilde *et al*., 2022). The constitutive activity of viral GPCRs modulates host gene expression, cell signaling events and host immunity, further contributing to the survival and proliferation of the virus (Coscoy, 2007). Previous studies have investigated the downstream oncogenic signaling pathways constitutively activated by KSHV-GPCR (Couty et al., 2001; Schwarz and Murphy, 2001; Shepard *et al*., 2001); however, the molecular and structural basis of the ligand-independent activation of this particular viral GPCR has yet to be unraveled. Recently, a study identified the structure of BILF1 in complex with human heterotrimeric Gi protein, providing a structural basis for ligand-independent BILF1-Gi signaling (Tsutsumi et al., 2021). Previous studies have also identified structures of a US27-Gi protein complex and a US28-CX3CL1-Gi protein complex (Tsutsumi et al., 2022). However, the signaling pathway downstream of US27 is still uncharacterized despite forming a complex with Gi proteins. Moreover, neither US27 nor BILF1 has identified endogenous ligand, whereas KSHV-GPCR binds the ELR-positive CXC chemokines CXCL1 and CXCL2 (Gershengorn et al., 1998). These chemokines further activate KSHV-GPCR above its constitutive levels, making KSHV-GPCR a structural and functional homolog of CXCR1 and CXCR2 (Gershengorn *et al*., 1998). Here, we present the cryo-EM structure of KSHV-GPCR-Gi protein complex at an overall resolution of 2.81 Å, and the cryo-EM structure of monomeric CXCL1-bound KSHV-GPCR-Gi protein complex at an overall resolution of 3.01 Å. Functional analysis confirmed that extracellular loop 2 (ECL2) and the structural motifs of activated GPCRs jointly play a crucial role in constitutive activation of KSHV-GPCR. With a nearly complete N terminal sequence of KSHV-GPCR solved (5 amino acids to the first Met), the cryo-EM structure of the CXCL1-bound KSHV-GPCR-Gi complex provides a useful model for detailed analysis of receptor interaction with CXCL1, which was not previously available.

## Methods

### Generation of recombinant CXCL1

The coding sequence of human CXCL1 with a C-terminal His6-tag was modified with a glycoprotein 67 (gp67) signal peptide. The CXCL1 construct was cloned into a pFastbac1 vector (ThermoFisher) and expressed in *Sf*9 insect cells as secreted proteins using baculovirus infection system. The media were collected after infection for 48 h. The pH of the supernatant was balanced by adding 1 M HEPES solution (pH 7.4). 1 mM nickel chloride and 5 mM calcium chloride were added and stirred for 1 h at 4 °C. The resulting precipitates were removed by centrifugation at 12,000 g and the supernatant was loaded onto Ni-NTA and incubated overnight. The nickel resin was washed with buffer containing 20 mM HEPES pH 7.4, 100 mM NaCl and 10 mM imidazole for 10 column volumes and then eluted in the above buffer containing 300 mM imidazole. The eluted CXCL1 were concentrated and purified over a size-exclusion chromatography column using a Superdex 75 column (GE Healthcare Life Sciences, Sweden). CXCL1 peak fractions were pooled, concentrated and fast-frozen by liquid nitrogen and stored at −80 °C for further usage.

### Preparation of apo-KSHV-GPCR-Gi and CXCL1-KSHV-GPCR-Gi complexes

The full-length KSHV-GPCR was used to obtain apo and CXCL1-bound signaling complexes. The N termini of KSHV-GPCR coding sequence were modified with an haemagglutinin peptide sequence, followed by a FLAG-tag, a human rhinovirus 3C (HRV 3C) protease cleavage site, and a thermostabilized apocytochrome b562RIL (BRIL) as a fusion protein to increase the protein expression and stability. A dominant-negative Gαi1 (DNGαi1) with two mutations (G203A and A326S) was generated by site-directed mutagenesis to decrease the affinity of nucleotide binding and limit G protein dissociation for stable complex. The DNGαi1 was cloned into a pFastbac1 vector and Gβ1γ2 was cloned into a pFastBac-Dual vector, respectively.

For expression of apo-KSHV-GPCR-Gi complex, the KSHV-GPCR, DNGαi1, Gβ1γ2 were co-expressed in Sf9 insect cells at a ratio of 1:1:1 when the cell density reached 2 × 10^6^ cells per ml. After 48 h of infection, the cells were collected by centrifugation at 2,000g for 20 min and the pellets were stored at −80°C for further purification. For expression of CXCL1-KSHV-GPCR-Gi complex, the baculovirus of CXCL1 was also added as the same ratio as KSHV-GPCR to facilitate the signaling complex formation.

For the purification of the apo-KSHV-GPCR-Gi complex, cell pellets from 2 L of culture were thawed at room temperature and suspended in the buffer containing 20 mM HEPES pH 7.4, 50 mM NaCl, 5 mM CaCl_2_ and 5 mM MgCl_2_ with 100× concentrated EDTA-free protease inhibitor cocktail (Roche, pallet). The suspensions were treated with Dounce and added with 25 mU ml^−1^ apyrase, followed by incubation for 1.5 h at room temperature. After incubation, the complex was extracted from the membrane with 0.8% (w/v) lauryl maltose neopentylglycol (LMNG; Anatrace, CAS. No. 1423315-02-8) and 0.1% (w/v) cholesteryl hemisuccinate (CHS; Anatrace, CAS. No. 102601-49-0) for 3 h at 4 °C. The supernatant was further isolated by centrifugation at 50,000g for 45 min and then incubated with pre-equilibrated Flag resin (20 mM HEPES pH 7.4, 100 mM NaCl) overnight at 4 °C. The Flag resin was loaded onto a gravity column manually. The resin was first washed with 15 column volumes of 20 mM HEPES, pH 7.4, 100 mM NaCl, 2 mM MgCl_2_, mM CaCl_2_, 0.01% LMNG (w/v), 0.002% CHS (w/v), and then eluted with the same buffer with 0.2 mg ml^−1^ FLAG peptide added. The eluted protein was concentrated to 500 μl with a 100-kDa molecular mass cut-off concentrator (Millipore). Concentrated apo-KSHV-GPCR-Gi complex was loaded onto a Superdex 200 Increase 10/300 GL column (GE Healthcare) with running buffer containing 20 mM HEPES pH 7.4, 100 mM NaCl, 0.00075% LMNG, 0.0002% CHS and 0.00025% GDN. The fractions for the monomeric complex were collected, evaluated by SDS-PAGE and concentrated to 11.4 mg ml^−1^ for cryo-EM experiments. For the CXCL1-KSHV-GPCR-Gi complex, the same steps were performed with adding purified CXCL1 during the process of protein purification, and the final sample was concentrated to 16 mg/ml for cryo-EM experiments.

### Cryo-EM grid preparation and data collection

For the cryo-EM grid preparation of the apo-KSHV-GPCR-Gi and CXCL1-KSHV-GPCR-Gi complexes, the 300 mesh Au R1.2/1.3 grids (Quantifoil) were used. The EM grids were glow-discharged by a Tergeo-EM plasma cleaner. Subsequently, 4ul of prepared protein sample was added to the grids. Then, the treated grids were blotted for 3 s and vitrified by plunging into liquid ethane in a Vitrobot Mark IV (Thermo Fisher Scientific) with 100% humidity at 4 °C. Finally, grids were stored in liquid nitrogen for further screening and data collection.

The Cryo-EM movie stacks were collected on an FEI Titan Krios microscope operated at 300 kV. The microscope was equipped with a Gatan Quantum energy filter. The movie stacks were collected using a Gatan K3 direct electron detector with a nominal magnification of ×105,000 in counting mode, resulting a pixel size of 0.85 Å. The energy filter was operated with a slit width of 20 eV. Each movie stack was dose-fractionated in 50 frames with a total exposure time of 2.5 s. The dose-rate was set at 20.8 e/pixel/s and the data was collected with a defocus ranging from −0.8 to −2.5 μm. Data collection was performed using SerialEM with one exposure per hole on the grid squares.

### Data processing and three-dimensional reconstruction

Movie stacks were subjected to beam-induced motion correction using MotionCor 2.1 The aligned micrographs were imported to cryoSPARC and further process was performed by cryoSPARC version v4.2.1. The Contrast transfer function (CTF) parameters for each micrograph were calculated by patch CTF estimation. The micrographs were manually inspected and obvious bad images were abandoned.

For the dataset of the apo-KSHV-GPCR-Gi complex, initial particle data-set was obtained by manual particle-picking. Then, 2D classification applied to generated template for automatic particle picking. Template-based particle selection yielded 2,695,689 particles that were subjected to reference-free two-dimensional (2D) classifications to discard bad particles. These particles were subsequently subjected to 3D classifications, 3D refinement, CTF refinement, local refinement, and generated a density map with an estimated global resolution of 2.81 Å at a Fourier shell correlation of 0.143.

For the datasets of CXCL1-KSHV-GPCR-Gi complexes, 2D templates for particle-picking was created by manual picking and 2D classification. Template-based particle selection yielded 2,545,563 particles that were subjected to rounds of reference-free 2D classifications to remove bad particles. After five rounds of maximum-likelihood-based 3D classifications, further 3D classification focusing on the receptor was performed. 107,180 particles from good classes were subsequently subjected to 3D refinement. After CTF refinement and local refinement, a final density map was obtained with an indicated global resolution of 3.01 Å at a Fourier shell correlation of 0.143.

### Model building, structure refinement and figure preparation

The models were manually built in COOT-0.9.8. Real-space refinements were performed using Phenix. The model statistics were validated using MolProbity. Structural figures were prepared in Chimera and PyMOL (https://pymol.org/2/). The final refinement statistics are provided in Supplementary Table S1. The maximum distance cut-off for polar hydrogen bond interactions and hydrophobic interactions were set at 3.5 Å and 4.5 Å, respectively.

### Mutagenesis study

Wild-type and mutant KSHV-GPCR were cloned into the pcDNA3.1(+) vector (Invitrogen). The coding sequences were chemically synthesized (General Biol) and cloned by overlap PCR using the ClonExpress Ultra One Step Cloning Kit (Vazyme Biotech; C115). All constructs were verified by DNA sequencing (Genewiz).

### cAMP inhibition assay

HeLa cells were transfected with receptor plasmids using Lipofectamine 3000 (Invitrogen) for 24 hrs. For the constitutive-active KSHV-GPCR model, different plasmid DNA concentrations were applied for transfection. On the following day, cells were harvested and washed twice using HBSS. Then cells were resuspended in the stimulation buffering containing HBSS with 5 mM HEPES, 0.1% BSA (w/v) and 0.5 mM 3-isobutyl-1-methylxanthine (IBMX). Different concentrations of CXCL1 (P6488; Beyotime, Shanghai, China) were prepared in the stimulation buffer. Cells were treated with the ligand at different concentrations (0 for the constitutive-active model) supplied with forskolin at a final concentration of 2.5 μM, then incubated for 30 mins at 37 °C. Intracellular levels of cAMP were detected using the LANCE Ultra cAMP kit (#TRF0263; PerkinElmer Life Sciences, Waltham, MA) following the manufacturer’s instructions. An Envision 2105 multimode plate reader (PerkinElmer) was used to collect the time resolved-fluorescence resonance energy transfer (TR-FRET) signals of intracellular cAMP levels.

### NanoBiT-based G protein dissociation assay

The dissociation of heterotrimeric G protein was examined using a NanoBiT-based G protein dissociation assay. HEK293T cells were plated in a 24-well plate for 24 hrs. Cells were transfected with a mixture of 100 ng pcDNA3.1(+) vector encoding KSHV-GPCR (WT/mutants), 100 ng pcDNA3.1 vector encoding Gαi1-LgBiT, 200 ng pcDNA3.1 vector encoding Gβ1 and 200 ng pcDNA3.1 vector encoding SmBiT-Gγ2 (per well in a 24-well plate), respectively. After 24 hrs transient expression, cells were collected and resuspended in 20 mM HEPES HBSS. Cells were then loaded onto a 384-well white plate (5,000 cells/well) and incubated with 10 μM coelenterazine H for 2 hrs at room temperature (Yeasen Biotech, Shanghai, China). Baseline chemiluminescence signals were read by an Envision 2105 multimode plate reader (PerkinElmer). CXCL1 at different concentrations was then added (0 for the constitutive-active model). For the constitutive-active model, raw baseline chemiluminescence signals were compared. For the ligand-dependent model, signals were measured after 15 mins’ incubation and normalized by the baseline readout. The percentage changes of signals were further calculated and plotted as a function of different concentrations of the chemokine ligand.

### Surface expression analysis

Since there is no commercially available flow cytometry antibody for KSHV-GPCR, a FLAG tag was added to the N-terminus of the receptor for flow cytometry analysis. Cells were transfected with FLAG-tagged WT or mutant KSHV-GPCR expression vectors for 24 hrs at 37°C. Then, cells were harvested and washed twice in HBSS containing 5% BSA on ice. Cells were then incubated with an Alexa Fluor™ 488-labeled anti-FLAG antibody (L5; Invitrogen, Cat # MA1-142; 1:50 diluted by HBSS buffer) for 30 mins on ice. The fluorescence signals for the antibody-receptor complex on the cell surface were then quantified by flow cytometry (CytoFLEX, Beckman Coulter, Brea, CA). Data were analyzed using FlowJo™ v10.8 Software (BD Life Sciences). Relative expression levels were represented according to the fluorescence signals measured in three independent experiments.

### Statistical analysis

For all cellular assays, at least three independent experiments were performed in duplicates. The data were expressed as mean ± standard error of means (SEM), and data analysis was performed in GraphPad Prism version 9.5.0. Analysis of Variance (ANOVA) using the one-way method was applied for statistical comparison. A *p*-value of 0.05 or lower is considered statistically significant.

## Results

### Overall structures of KSHV-GPCR-Gi complex without and with bound CXCL1

Stable complexes of KSHV-GPCR-Gi protein were reconstituted by co-expression of KSHV-GPCR and heterotrimeric Gi proteins in *Sf9* insect cells (Fig. S1A). The purified receptor-Gi protein complexes were then subjected to single-particle cryo-EM analysis, yielding a structural model of the KSHV-GPCR-Gi complex at a global resolution of 2.81 Å (Fig. 1 A, 1B; Fig. S1 B-F). The monomeric CXCL1-bound KSHV-GPCR-Gi complex was reconstituted by co-expression of human CXCL1 together with the receptor and Gi proteins in Sf9 cells. Single-particle analysis of the CXCL1-KSHV-GPCR-Gi complex produced a structural model at a global resolution of 3.01 Å (Fig. 1C and 1D, Fig. S2 B-F). The reconstituted 3D structures revealed a typical seven-transmembrane (7-TM) topology of GPCRs with either ligand-free (Fig. 1B) or CXCL1-bound (Fig. 1D) TM pocket. The overall structures of CXCL1-bound and constitutively-active KSHV-GPCR are nearly identical, with a root mean square distance (RMSD) of 0.32 Å between their C⍺ atoms (CXCL1 excluded). In both structures, the disulfide bond between ECL1 (C118^ECL1^) and ECL2 (C196^ECL2^) is clearly visible, whereas the predicted disulfide bond between N terminus (C39^NT^) and ECL3 (C286^ECL3^) is not. A close examination found that the two cysteines were 3.6 Å apart, making it difficult to form a disulfide bond. The N terminal Pro-Cys (PC motif) found in most chemokine receptors becomes Val-Cys in KSHV-GPCR and lacks the bent conformation. The constitutively active KSHV-GPCR interacts extensively with the α5 helix of Gαi, and to a close proximity with the αN helix of Gαi and Gβ subunit, thereby securing the conformation of an active receptor-Gi protein complex as is commonly seen in agonist-bound Class A GPCRs (Weis and Kobilka, 2018). To verify the ligand-independent constitutive activity of KSHV-GPCR, we further examined Gi signaling of the receptor by analyzing basal levels of cAMP accumulation and G protein dissociation without ligand. In both assays, cAMP inhibition and G protein dissociation were correlated positively with the amount of input plasmid DNA coding for KSHV-GPCR (Fig. 1 E and F).

**Figure 1.**
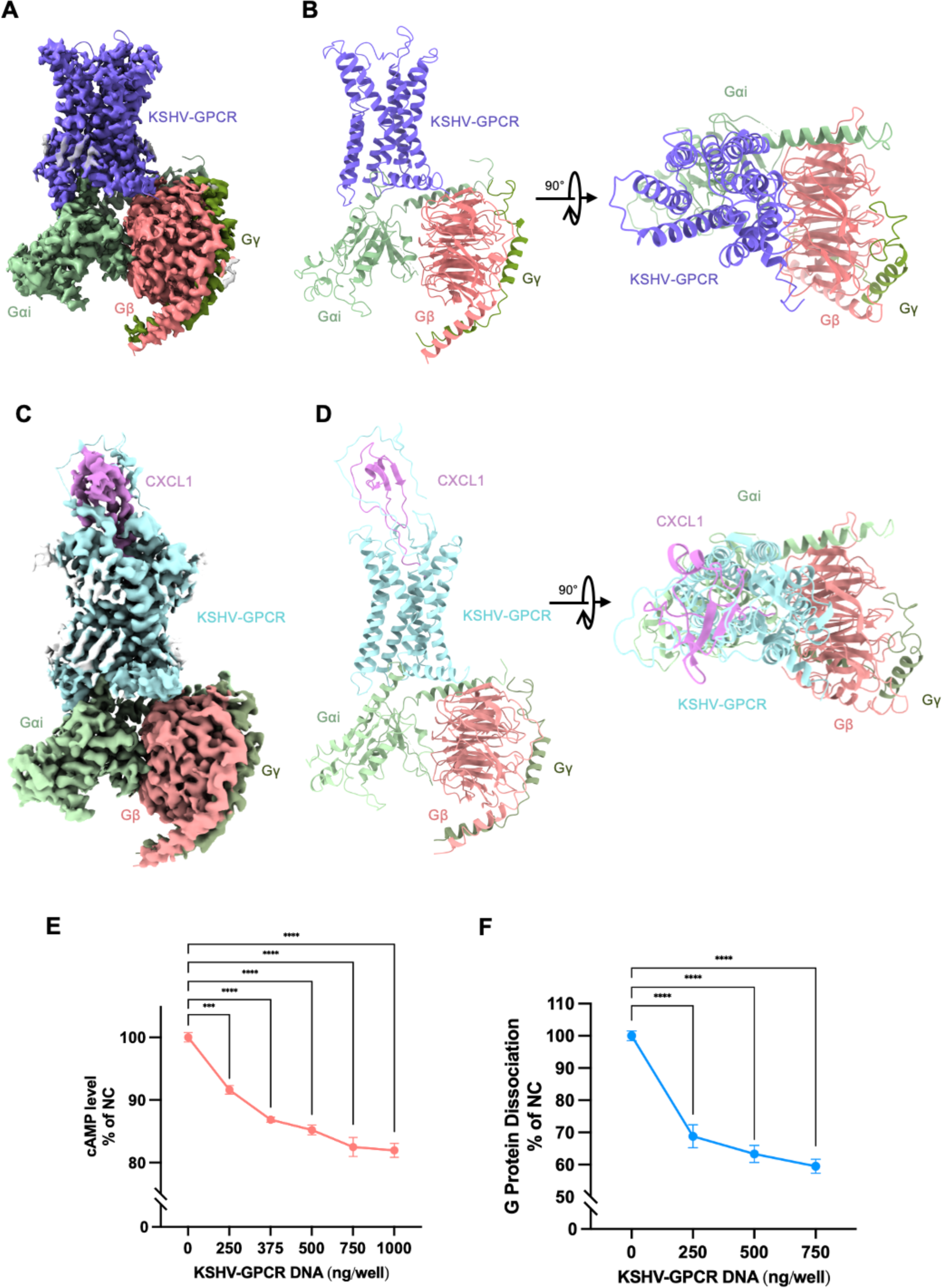
Overall structure of KSHV-GPCR-Gi complex and KSHV-GPCR-CXCL1-Gi complex. **A** Cryo-EM electron density map of constitutive-active KSHV-GPCR in complex with heterotrimeric Gi proteins at an overall resolution of 2.81 Å. Ligand-free constitutive-active KSHV-GPCR is shown in purple, Gαi in lime green, Gβ in salmon red and Gγ in dark green. **B** Atomic coordinates of constitutive-active KSHV-GPCR in complex with heterotrimeric Gi proteins from side view (left panel) and top view (right panel) Ligand-free constitutive-active KSHV-GPCR is shown in purple, Gαi in lime green, Gβ in salmon red and Gγ in dark green. **C** Cryo-EM electron density map of CXCL1-bound KSHV-GPCR in complex with heterotrimeric Gi proteins at an overall resolution of 3.01 Å. CXCL1-bound KSHV-GPCR is shown in cyan, CXCL1 in magenta, Gαi in lime green, Gβ in salmon red and Gγ in dark green. **D** Atomic coordinates of CXCL1-bound KSHV-GPCR in complex with heterotrimeric Gi proteins from side view (left panel) and top view (right panel). CXCL1-bound KSHV-GPCR is shown in cyan, CXCL1 in magenta, Gαi in lime green, Gβ in salmon red and Gγ in dark green. **E** Basal levels of cAMP accumulation without ligand treatment. **F** Basal levels of NanoBiT-based G protein dissociation without ligand treatment. Data were obtained from three independent experiments, each with three replicates.

### Extracellular Loop 2 is crucial for constitutive activation of KSHV-GPCR

Constitutively active GPCRs display high basal activity without agonist binding. Previous reports illustrated the possibility that a large extracellular loop 2 (ECL2) may provide additional contacts for constitutive activation of a viral chemokine receptor (Tsutsumi *et al*., 2021). The ECL2 in KSHV-GPCR (a.a. 183-207) is the largest among the 3 extracellular loops (Fig. 2A, 2B), and its role in constitutive activation of the receptor was examined by substitution of the loop with ECL2 counterparts from several other Class A GPCRs including CXCR2, FPR1, CXCR4 and CMKLR1 (Fig. 2 C-F). These mutants were further subjected to functional analysis using NanoBiT-based complementation assay (Dixon et al., 2016) for measurement of G protein dissociation. Basal levels of G protein dissociation were compared among different mutants and with the wild-type KSHV-GPCR control. Significantly elevated chemiluminescence signals were observed for ECL2 substituted mutants, indicating that constitutive activation of G protein was abrogated in the ECL2 substituents. These findings further suggested an important role of ECL2 in the constitutive activation of KSHV-GPCR.

**Figure 2.**
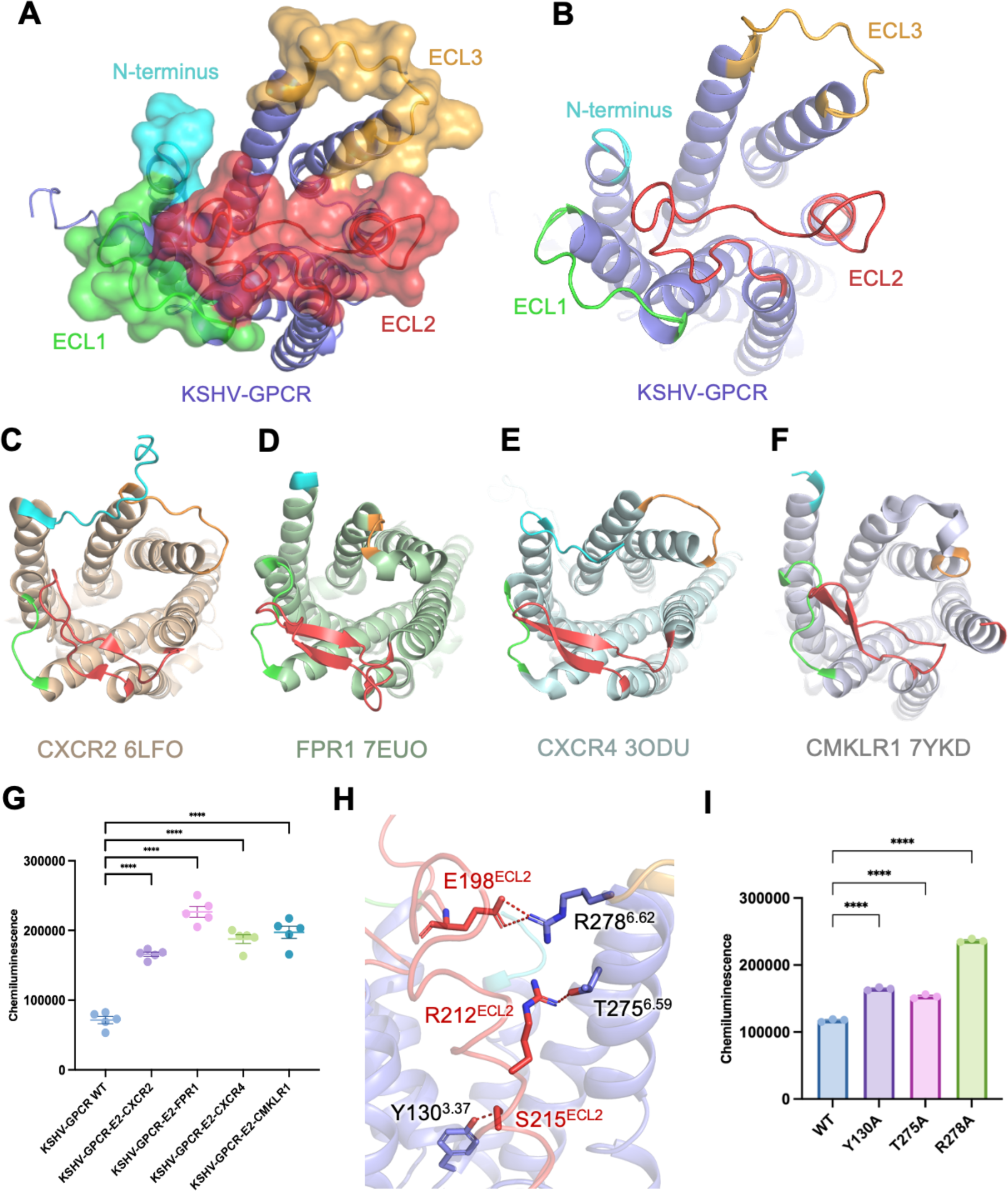
Extracellular loop 2 of KSHV-GPCR is crucial for constitutive G protein activation. **A** The structure of constitutive-active KSHV-GPCR in an extracellular view. Receptor N terminus and extracellular loops are shown as surface. N terminus loop is colored in cyan, ECL1 is marked with green, ECL2 is shown in red and ECL3 is highlighted in yellow. **B** The structure of constitutive-active KSHV-GPCR in an extracellular view. Receptor N terminus and extracellular loops are shown as cartoon. ECL2 of KSHV-GPCR occupies a relatively large space over the canonical binding pocket of the receptor. **C-F** Extracellular view of the structures of **(C)** CXCR2 (active, PDB ID: 6LFO), **(D)** FPR1 (active, PDB ID: 7EUO), **(E)** CXCR4 (inactive, PDB ID: 3ODU), and **(F)** CMKLR1 (PDB ID: 7YKD). **G** ECL2 of KSHV-GPCR was substituted by ECL2 of CXCR2, FPR1, CXCR4 and CMKLR1, respectively. Basel levels of G protein dissociation were tested and compared among mutants and wildtype control. **H** Polar interaction between ECL2 of KSHV-GPCR and the canonical transmembrane binding pocket. **I** Reduced ligand-independent constitutive G protein activation in mutants.

To investigate how ECL2 engages in the constitutive activation of KSHV-GPCR, we focused on polar contacts between ECL2 and the rest of the receptor. Among the amino acid residues examined in the constitutively active KSHV-GPCR model, E198^ECL2^ forms multiple polar bonds with R278^6.62^. Additionally, R212^ECL2^ interacts with T275^6.59^, and S215^ECL2^ forms polar interaction with Y130^3.37^ (Fig. 2H). Alanine substitutions of Y130^3.37^, T275^6.59^ and R278^6.62^ were performed and the resulting mutants were subjected to a NanoBiT-based G protein dissociation assay. As shown in Fig. 2I, significantly elevated chemiluminescence signals were observed with all 3 mutants, demonstrating reduced G protein activation. These mutants readily expressed on cell surfaces (Fig. S4). Altogether, these results imply a direct engagement of ECL2 in constitutive activation of KSHV-GPCR.

### Recognition of CXCL1 by KSHV-GPCR

Next, we examined the cryo-EM structure of the CXCL1-bound KSHV-GPCR-Gi complex. CXCL1 is one of the full agonists of KSHV-GPCR and further activates the receptor above constitutive level (Gershengorn *et al*., 1998). In the CXCL1-KSHV-GPCR-Gi complex, CXCL1 was solved from A1 to I61 following the first turn of the C-terminal helix, after which there was insufficient electron density for modeling (Fig. 3A). As an ELR-positive chemokine, E6-L7-R8 presents a ELR motif in CXCL1. The CXC motif (C9-Q10-C11) in the N-loop forms two disulfide bonds for connections to the 30s-loop and the β3-strand (Fig. 3A). The globular core of CXCL1 is defined by β1-strand, 30s-loop, β2-strand, 40s-loop, β3-strand and a C-terminal helix. In the CXCL1-KSHV-GPCR structure, the chemokine and the receptor are in 1:1 stoichiometry, and their interaction appears to follow the two-site model (Scholten et al., 2012; Urvas and Kellenberger, 2023) with clearly identifiable chemokine recognition sites 1, 2 and 1.5 (CSR1, CSR2 and CSR1.5) (Fig. 3B). At CSR1, the globular core of CXCL1 interacts with the N terminus of KSHV-GPCR, which was solved to the 5^th^ amino acid. Notably, the N terminus of KSHV-GPCR fits into a groove on CXCL1 in between the β1-strand and the C-terminal helix (Fig. 3C). Extensive hydrophobic contacts between the residue pairs are observed at CRS1, including D5^NT^-S30, F6^NT^-V28, L7^NT^-V28, T8^NT^-N27, I9^NT^-V26, L11^NT^-I23, W17^NT^-K21 (Fig. 3C). In addition, multiple polar interactions are present between D13^NT^ of the receptor and K21 of CXCL1, and between Y26^NT^ of the receptor and CXCL1 K61 (Fig. 3D). These interactions allow the N terminus of KSHV-GPCR to wrap around the chemokine core tightly.

**Figure 3.**
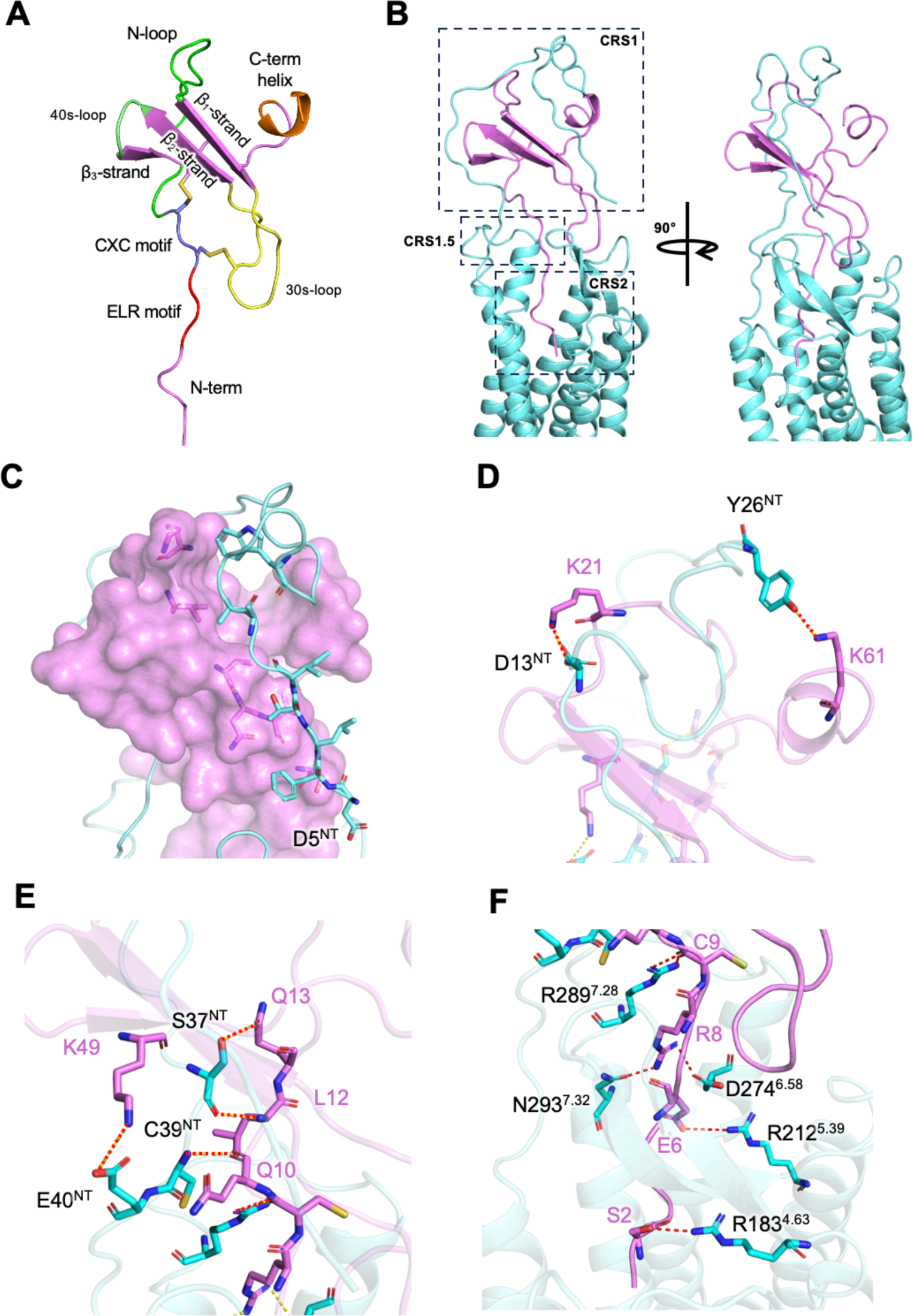
Molecular interactions between CXCL1 and KSHV-GPCR. **A** Structure of CXCL1. Structural regions of CXCL1 are highlighted, including N-terminus (magenta), ELR motif (red), CXC motif (slate blue), N-loop(green), 30s-loop (yellow), 40s-loop (grass green), β1-β3 strands (magenta), and C-terminal helix (orange). **B** Overview of the interaction between CXCL1 (magenta) and KSHV-GPCR (cyan). Chemokine binding sites CRS1, CRS1.5, and CRS2 are marked with dashed square. **C** The N-terminus of KSHV-GPCR (resolved to the fifth amino acid D5) wraps around the globular core of CXCL1. Residues with hydrophobic interactions are shown in sticks. CXCL1 is shown in surface representation. **D** Polar contacts between CXCL1 and KSHV-GPCR. Hydrogen bonds (red dotted dashes) are observed between D13^NT^ and K21, Y26^NT^ and K61 (shown in sticks), respectively. **E** Polar interactions between CXCL1 and KSHV-GPCR at chemokine recognition site 1.5 (CRS1.5). **F** Polar interactions between CXCL1 and KSHV-GPCR at chemokine recognition site 2 (CRS2). Hydrogen bonds are highlighted in dotted lines.

At CRS1.5, defined by the PC motif of the receptor, extensive polar interactions hold together the receptor PC motif (VC in KSFV-GPCR) and the CXC motif of CXCL1. S37^NT^ forms two polar bonds with L12 and Q13 of CXCL1, C39^NT^ interacts with the backbone oxygen atom of Q10, and E40^NT^ interacts with K49 on the β3-strand of CXCL1 (Fig. 3E).

The N terminus of CXCL1 protrudes deep into the transmembrane binding pocket of the receptor that forms CRS2. Here, R289^7.28^ interacts with the backbone of C9. There are two polar contacts: D274^6.58^ and N293^7.32^ form polar interactions with R8 of CXCL1, R212^5.39^ interacts with E6, and R183^4.64^ at the bottom of the binding pocket forms polar interaction with S2 of CXCL1 (Fig. 3F). The ELR motif at chemokine N terminus contributes to the main intermolecular polar interactions at CRS2. For non-polar interactions, A1 of CXCL1 forms multiple hydrophobic contacts with L54^1.38^, E103^2.60^, E121^3.27^. A4 has non-polar interactions with A297^7.36^. T5 forms non-polar interactions with Y197^ECL2^. Moreover, the N terminus of CXCL1 favors the minor sub-pocket of KSHV-GPCR in proximity to TM1-3.

Functional validation of receptor-chemokine interactions was further conducted through NanoBiT-based G protein dissociation assay and site-directed mutagenesis. Alanine substitution at CRS1 (D13^NT^A and Y26^NT^A) did not completely eliminate the G protein response (Fig. 4A). At CRS1.5, alanine substitutions including S37A, C39A and E40A markedly diminished G protein activation upon CXCL1 stimulation (Fig. 4B) while having little or no effect on the constitutive activity of KSHV-GPCR. Deep into the TM pocket, alanine substitution at CRS2 (R183A, R212A, D274A, R289A, and N293A) abrogated the G protein dissociation response (Fig. 4C, 4D). Cell surface expression of the mutants was comparable to WT (Fig. S4). These results confirmed the two-state model of receptor-chemokine interaction, in which CRS1 is responsible for the initial recruitment of chemokines but does not alter the downstream G protein signaling, while CRS1.5 and CRS2 contribute more to the potency and efficacy of chemokine activation of G proteins.

**Figure 4.**
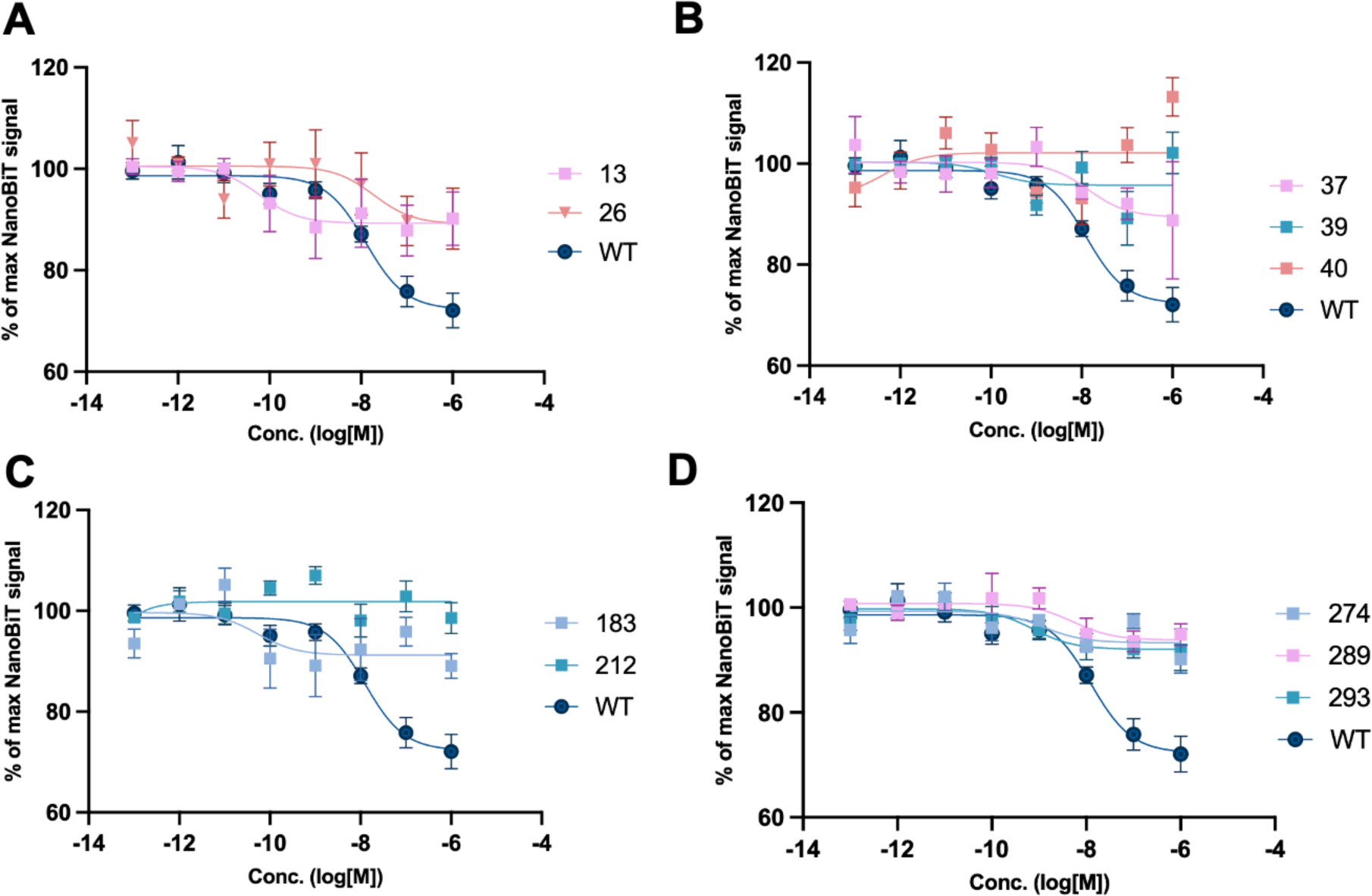
Functional analysis of CXCL1 binding to KSHV-GPCR. Alanine substitution was conducted for CXCL1-binding residues of KSHV-GPCR. NanoBiT-based G protein dissociation assay was performed in cells transiently expressing wildtype or mutated KSHV-GPCR to evaluate the importance of each residue to CXCL1 binding. **(A)** CRS1 mutants retained some G protein activity, while mutations at **(B)** CRS1.5 and **(C-D)** CRS2 completely abolished G protein activation upon CXCL1 treatment.

### Comparison of chemokine binding mode between KSHV-GPCR and CXCR2

The structures of CXCL1-bound and constitutive-active KSHV-GPCR are nearly identical, but small differences are observed adjacent to chemokine binding sites (Fig. 5A). To accommodate the binding of CXCL1, ECL2 of KSHV-GPCR tilts upwards, thereby disengaging the interactions with the binding cavity. TM3 bends towards the center of the transmembrane binding pocket (Fig. 5A, left). From an extracellular view of the receptor, it is well-observed that the β-hairpin region of ECL2 and ECL1 move closer to the chemokine for accommodation of the chemokine core (Fig. 5A, left). Given the sequence homology of KSHV-GPCR and CXCR2, it is speculated that KSHV-GPCR is hijacked from CXCR2-expressing host cells by KSHV (Rosenkilde *et al*., 2022). We therefore further compared the structures of CXCL1-bound KSHV-GPCR and CXCL8-bound CXCR2 (Fig. 5B, 5C). Compared with CXCL8-CXCR2 interactions, the N-terminus of CXCL1 protrudes 7.9 Å deeper into KSHV-GPCR receptor binding cavity at CRS2 (Fig. 5B, left), and the globular core of CXCL1 rotates ∼40° counter-clockwise from an extracellular point of view (Fig. 5B, right). A series of counter-clockwise inward bends are observed at the extracellular regions of TM1, 2, and 3. The top of TM loop 4 bends slightly outward, and TM7 displays an inward movement to the center of the chemokine binding pocket (Fig. 5C).

**Figure 5.**
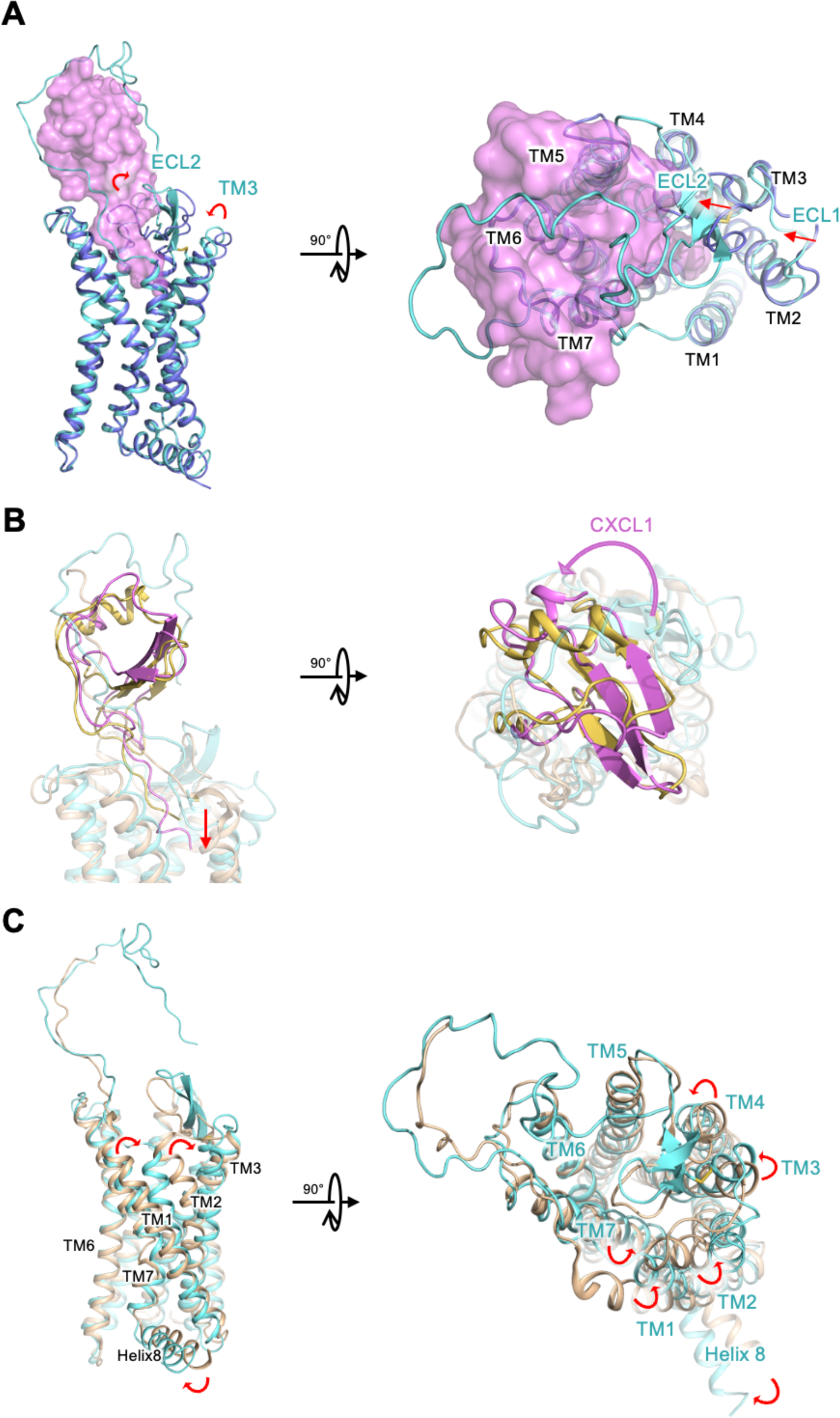
Comparison between CXCL1-bound KSHV-GPCR and constitutive active KSHV-GPCR, and CXCL1-bound KSHV-GPCR and CXCL8-bound CXCR2. **A** An inward movement of receptor ECL1, ECL2 and TM3 was observed for CXCL1-bound KSHV-GPCR (cyan), compared with the ligand-free constitutive active structure of KSHV-GPCR (purple). **B** Comparison of chemokine orientations between CXCL1-bound KSHV-GPCR (CXCL1: magenta, KSHV-GPCR: cyan) and CXCL8-bound CXCR2 (CXCL8: yellow, CXCR2: wheat). CXCL1 inserts deeper into the CRS1 of KSHV-GPCR (left). The globular core of CXCL1 shifts towards ECL3 (right). **C** Comparison of receptor conformations between CXCL1-bound KSHV-GPCR (CXCL1: magenta, KSHV-GPCR: cyan) and CXCL8-bound CXCR2 (CXCL8: yellow, CXCR2: wheat). At TM1, TM2, TM3, TM4 and TM7, inward orientation of helices were observed for CXCL1-bound KSHV-GPCR.

### Molecular switches of KSHV-GPCR and their roles in ligand-independent activation

To answer the question that how KSHV-GPCR has structurally evolved from its host analog into a constitutively active receptor, we examined the molecular switches conserved among class A GPCRs. The structure of KSHV-GPCR was aligned with that of CXCR2 in either active state (Fig. 5B) or inactive state (Fig. 6 A and B), and the structural motifs N^7.49^P^7.50^xxY^7.53^, CW^6.48^xP^6.50^, P^5.50^-I^3.40^-F^6.44^ were compared. In the NPxxY motif, N^7.49^ is substituted by valine in KSHV-GPCR (expressed as V/N^7,49^). In CW^6.48^xP^6.50^, the highly conserved W^6.48^ is substituted by a cysteine (C/W^6,48^) that greatly alters the spatial occupancy. Since substitution at these conserved motifs may create hyperactive or constitutive active class A GPCRs (Burger et al., 1999; Fritze et al., 2003), we further looked into the details of the geometry of residues lining these structural motifs (Fig. 6 C-H).

**Figure 6.**
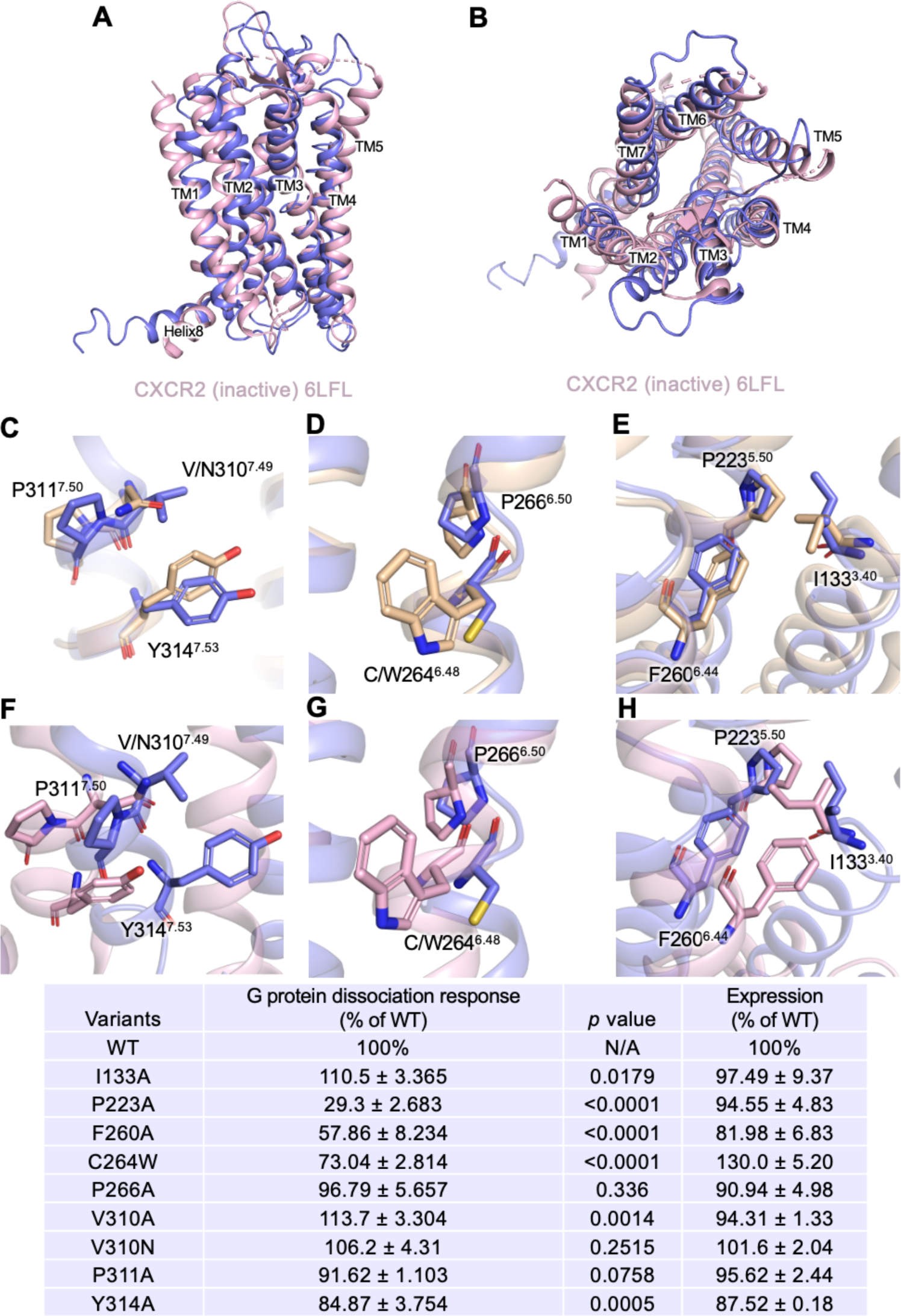
Comparison of conserved structural motifs between active KSHV-GPCR and either active or inactive CXCR2. **A** Side view of superimposed KSHV-GPCR (purple) and inactive CXCR2 (light pink). **B** Top view of superimposed KSHV-GPCR (purple) and inactive CXCR2 (light pink). **C-E** Detailed comparison between KSHV-GPCR (purple) and active CXCR2 (wheat gold). **(C)** NPxxY motif, **(D)** P^6.50^-C/W^6.48^ “toggle switch”, and **(E)** P^5.50^-I^3.40^-F^6.44^ motif were compared. **F-H** Detailed comparison between KSHV-GPCR (purple) and inactive CXCR2 (light pink). **(F)** NPxxY motif, **(G)** P^6.50^-C/W^6.48^ “toggle switch”, and **(H)** P^5.50^-I^3.40^-F^6.44^ motif were compared. **Table** Basel levels of NanoBiT signals in G protein dissociation assay. Residues of these structural motifs were substituted by alanine. Readouts were normalized and shown as percentages of G protein response of WT strain.

N^7.49^P^7.50^xxY^7.53^ motif in class A GPCR mediates the transition from inactive to active conformation for G protein binding. However, the mutation of V^7.49^ back to asparagine did not markedly change the constitutive activity of the receptor. Alanine substitution was conducted at V^7.49^, P^7.50^ and Y^7.53^, respectively. No significant change in the extent of constitutive G protein dissociation was observed, suggesting that other players may also participate in the activation process.

Another structural motif, CW^6.48^xP^6.50^ is mutated to CC^6.48^xP^6.50^. This “toggle switch” motif works as a rotamer switch, bending the cytoplasmic half of TM6 outward to accommodate the binding of G protein. By converting to cysteine at this position, the residue with a smaller side chain can rearrange the orientation of TM6 for activation (Fig. 6 D and G). Indeed, by mutating C^6.48^ back to tryptophan, the constitutive G protein activation was reduced significantly. This implies the importance of C^6.48^ in constitutive G protein activation.

We next analyzed the P^5.50^-I^3.40^-F^6.44^ motif, which forms an interface between TM3, TM5 and TM6. The conformational change of this motif is responsible for the outward displacement of TM6. The orientations of residue side chains in this motif are nearly identical to those in an active CXCR2, favoring G protein activation (Fig. 6 E and H). Alanine substitution at P^5.50^ and F^6.44^ abrogated the constitutive G protein dissociation, but alanine substitution of I^3.40^ did not significantly affect the constitutive activation of KSHV-GPCR (p = 0.0179).

Altogether, based on structural geometry and signaling properties at these conserved molecular switch motifs, our KSHV-GPCR structural model supports an active conformation. Among these highly conserved residues, some substitutions found exclusively at KSHV-GPCR may contribute to its constitutive activation by reshaping the G protein binding cavity.

### Receptor-Gi interface of KSHV-GPCR compared with active and inactive CXCR2

Next, we examined the receptor-Gi interface of KSHV-GPCR. The DRY motif forms direct polar contact with the Gαi protein and is crucial for G protein activation. In KSHV-GPCR, D^3.49^ is substituted by valine. By reversely mutating valine at position 3.49 back to D, no significant deviation in the extent of G protein activation was observed. Polar interactions between the receptor and Gαi protein involve R^3.50^-C351, L^3.53^-N347, as well as Q^6.28^-E318 pairs (Fig. 7 B and D). Alanine substitution of R^3.50^ and L^3.53^ resulted in a loss of G protein activity, while for the Q^6.28^A mutant, no significant difference was observed. The ionic interactions in the G protein binding cavity are highly important for G protein activation. Despite the high overall similarity in structures between KSHV-GPCR and the active CXCR2, there are some differences at the receptor-Gi interfaces (Fig. 7 A and B). First, a displacement toward the cytoplasm is observed for TM3, resulting in a deeper protrusion of the α5 helix of the Gαi protein into the G protein binding cavity. Secondly, the intracellular loop 2 (ICL2) of KSHV-GPCR is closer to the Gi protein, while in active CXCR2, ICL2 bends outwards. By comparing the inactive CXCR2 structure with the KSHV-GPCR structure, several distinct features support the active state model of KSHV-GPCR (Fig. 7 C and D). In TM3, the geometry and position of the DRY motif are similar in both structures, yet R^3.50^ in KSHV-GPCR takes a more relaxed pose, with better contact with the α5 helix in the G protein binding cavity. The outward movement of TM6 of KSHV-GPCR in its active state allows direct contact between Q^6.28^ and Gαi protein, while the ICL3 and adjacent TM6 in the inactive CXCR2 structure is distant from the active conformation. These results indicate that the G protein-activating capacity of KSHV-GPCR is robust, given the orientations and geometry of its TM3 residues lining the G protein binding cavity.

**Figure 7.**
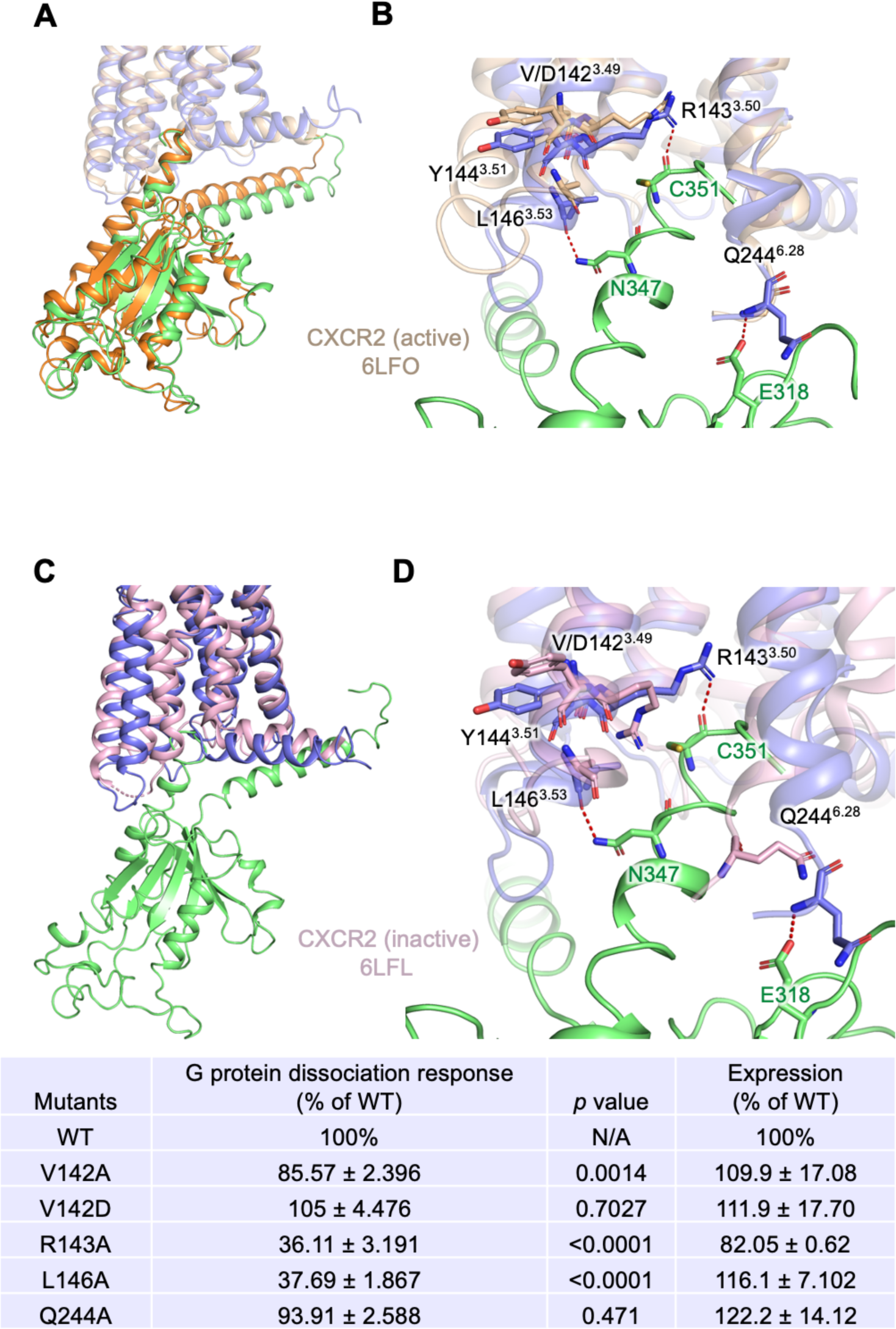
Comparison of receptor-Gαi interface between active KSHV-GPCR and either active or inactive CXCR2. **A** Side view of superimposed KSHV-GPCR (purple) and active CXCR2 (wheat gold) in complex with Gαi protein (green). **B** Close-up view of superimposed KSHV-GPCR (purple) and active CXCR2 (wheat gold) in complex with Gαi protein (green). Residues with polar interaction were highlighted in sticks. **C** Side view of superimposed KSHV-GPCR (purple) and inactive CXCR2 (light pink) in complex with Gαi protein (green). **D** Close-up view of superimposed KSHV-GPCR (purple) and inactive CXCR2 (light pink) in complex with Gαi protein (green). **Table** Basel levels of NanoBiT signals in G protein dissociation assay. Alanine substitution was conducted to residues at receptor-Gi interface of KSHV-GPCR. Readouts were normalized and shown as percentages of G protein response of WT strain.

## Discussion

In this work, we present cryo-EM structures of KSHV-GPCR in active conformation. KSHV-GPCR, encoded by ORF74, is expressed in the lytic cycle of HHV-8 and is responsible for angiogenesis and oncogenic transformation of Kaposi’s sarcoma (Bais *et al*., 1998). Exogenous expression of KSHV-GPCR in mice led to Kaposi’s sarcoma-like pathological changes (Cesarman *et al*., 2000), indicating a crucial function of the constitutively active receptor in oncogenic transformation. Based on our cryo-EM model of the unliganded KSHV-GPCR, ECL2 is essential for the constitutive activity. ECL2 folds back to the opening of the TM pocket and forms polar interactions with KSHV-GPCR through the E198^ECL2^-R278^6.62^, R212^ECL2^-T275^6.59^, and S215^ECL2^-Y130^3.37^ pairs. Disruption of these interactions with alanine substitutions of Y130^3.37^, T275^6.59^ and R278^6.62^, in particular the R278^6.62^ substitution, reduced the constitutive activity. Moreover, the substitution of ECL2 with the counterparts of CXCR2, FPR1, CXCR4 and CMKLR1, all depending on agonist binding for receptor activation, abrogated constitutive activation of the resulting chimeras. At the receptor-Gi protein interface, there is no significant change between the unliganded and CXCL1-bound KSHV-GPCR, suggesting that ECL2 is able to hold the receptor in an active conformation for Gi protein activation.

Constitutive activity was also observed in other viral GPCRs, notably US28 from human cytomegalovirus (HCMV) and BILF1 from Epstein Barr Virus (EBV). US28 has a TM pocket for binding of CC chemokines of the MIP and MCP classes, and of CX3CL1 in a different mode (Rosenkilde *et al*., 2022). Still, the two-site model applies to these interactions between the receptor and its various ligands. US28 displays constitutive activity through Gi, Gq and G12/13 pathways. However, the binding of CX3CL1 to US28 is context-dependent, as this ligand can serve as either a partial agonist or an inverse agonist (Burg et al., 2015). In contrast, BILF1 does not have an identified chemokine ligand, and its TM pocket is occluded by the ECL2 (Tsutsumi *et al*., 2021). This structural feature is responsible for the EBV oncogenic function of BILF1 by inducing constitutive signaling mainly through the Gi proteins (Tsutsumi *et al*., 2021). Like BILF1, KSHV-GPCR has an occluded orifice due to polar interactions between ECL2 and the side chain of TM3, TM5 and TM6.

However, KSHV-GPCR binds a number of CC and CXC chemokines that further activate the receptor. Based on sequence comparison, KSHV-GPCR has the highest homology with human CXCR2 (27% identical amino acids). Therefore, KSHV-GPCR is a CXCR hijacked by the virus. In this study, we chose CXCL1, also termed GRO-α, for its melanoma growth-stimulating activity and transforming capability through CXCR2. Our structural model shows a clearly defined TM pocket that accommodates CXCL1 and serves as CRS2. CXCL1 binding disrupts the polar interactions between ECL2 and the side chain of TM3, TM5 and TM6, but maintains the active conformation of KSHV-GPCR based on structural comparison of the unliganded and CXCL1-bound receptor. At CRS2, the ELR motif of CXCL1 is a critical determinant of the potency of the agonist by polar interactions between E6 and R212^5.39^. In addition, R8 interacts with both D274^6.58^ and N293^7.32^. Disruption of these important interactions at CRS2 abrogates the CXCL1-induced G protein dissociation, confirming that CRS2 is essential to the induced activation of KSHV-GPCR by the chemokine.

Before high-resolution chemokine receptor structures became available, domain swapping experiments were carried out and led to the prediction that the N terminus of KSHV-GPCR is necessary for binding of chemokine ligand but not for transmembrane signaling (Ho et al., 1999). The N terminus of KSHV-GPCR (1-47) contains a stretch of negatively charged amino acids, notably 12-DDDE-15. Moreover, the disulfide bond between the PC motif (VC in KSHV-GPCR) and C286^ECL3^ is not found in this structure. These features presumably make the N terminus of the receptor more flexible in the initial interaction with chemokines at CRS1. In the reconstructed 3D model of CXCL1-bound KSHV-GPCR-Gi complex, the receptor N terminus wraps around the globular core of CXCL1. Analysis of a fully settled CXCL1 in the TM pocket shows D13^NT^ interaction with K21 of CXCL1, and Y26^NT^ interaction with K61. These polar interactions serve to stabilize the bound chemokine. As a transition from CRS1 to CRS2, CRS1.5 contains several residues in the receptor N terminus that interact with the N-loop and β3-strand of CXCL1, including Q10, L12, Q13 of the N-loop and K49 of the β3-strand. These interactions may be helpful to guide the N terminus of the chemokine into the TM pocket, as indicated by data from our mutagenesis assay.

Early studies, conducted shortly after the discovery of KSHV-GPCR, focused on key residues that profoundly influence the constitutive activity of the receptor (Ho et al., 2001). These studies did not identify key residues unique to KSHV-GPCR. For example, the substitution of R143^3.50^, which is a part of the critical DRY motif, abolished all Class A GPCR signaling. This and other findings suggest that the constitutive activity of KSHV-GPCR is the combined outcome of mutations of the hijacked receptor gene in the virus (Burger *et al*., 1999; Ho *et al*., 2001).

Nevertheless, the “ionic lock” characterized by the interaction between D/E^3.49^, R^3.50^ and D/E^6.30^ stabilizes the inactive conformation of GPCRs (Ballesteros et al., 2001). In KSHV-GPCR, D/E^3.49^ and D/E^6.30^ are substituted by V^3.49^ and R^6.30^, respectively. As demonstrated by previous studies and in line with our mutagenesis data, V^3.49^D substitution did not decrease the constitutive activity of the receptor. Besides, the geometry of R^6.30^ is distant from V^3.49^ and R^3.50^, thereby disrupting the “ionic lock”. Another important and conserved residue, W^6.48^ for TM6 conformational change, is substituted by cysteine in KSHV-GPCR. This mutation may have a strong impact on KSHV-GPCR constitutive activity, as C^6.48^W disrupted the ligand-independent G protein activation. The canonical NPxxY motif is characterized by a hydrogen bond network between N^7.49^ and D^2.50^. However, these residues are mutated to V^7.49^ and S^2.50^ in KSHV-GPCR, respectively. The two residues are distant from each other, which is unable to support the canonical hydrogen bond network. As evidenced in our study and other biochemical studies, V^7.49^N and S^2.50^D mutations did not alter the constitutive activity. Previous study suggested the importance of L^2.48^ and L^2.51^ in constitutive activity, as the L^2.48^D and L^2.51^D mutants diminished the constitutive activity but could still be activated by chemokine ligands, and KS lesions were absent in transgenic mice carrying L^2.48^D mutant (Cesarman *et al*., 2000; Holst et al., 2001). In line with their hypothesis, L^2.48^ and L^2.51^ in our KSHV-GPCR structures indeed face the lipid bilayer, supporting the active conformation in both unliganded and CXCL1-bound KSHV-GPCR. Substitution by charged amino acids may destabilize the active conformation of the receptor, thus requiring extra energy (ligand binding) for effective conformational changes toward activation.

In conclusion, our cryo-EM model provides the first solved structure of a CXC chemokine bound to a viral GPCR that possesses both constitutive and ligand-induced activities. Moreover, this study also provides the first high-resolution structure of CXCL1 bound to a chemokine receptor. These structural information will likely shed light on how viral GPCRs maintain constitutive activity, which may help us to understand chemokine binding and activation mechanisms in general.

## Acknowledgments

This work was supported in part by grants from the Science, Technology and Innovation Commission of Shenzhen Municipality GXWD20201231105722002-20200831175432002 (R.D.Y.), and Shenzhen Science and Technology Program Grant No. RCBS20221008093330067 (A.L.). This work was also supported by National Natural Science Foundation of China 32070950 (R.D.Y.), China Postdoctoral Science Foundation 2022M713049 (A.L.), the Ganghong Young Scholar Development Fund (R.D.Y.), the fund from Kobilka Institute of Innovative Drug Discovery at The Chinese University of Hong Kong, Shenzhen (R.D.Y.), and the fund from Shenzhen-Hong Kong Cooperation Zone for Technology and Innovation (HZQB-KCZYB-2020056).

## Competing Interest Statement

The authors declare no competing interest.

## SUPPLEMENTARY INFORMATION

**Fig. S1.**
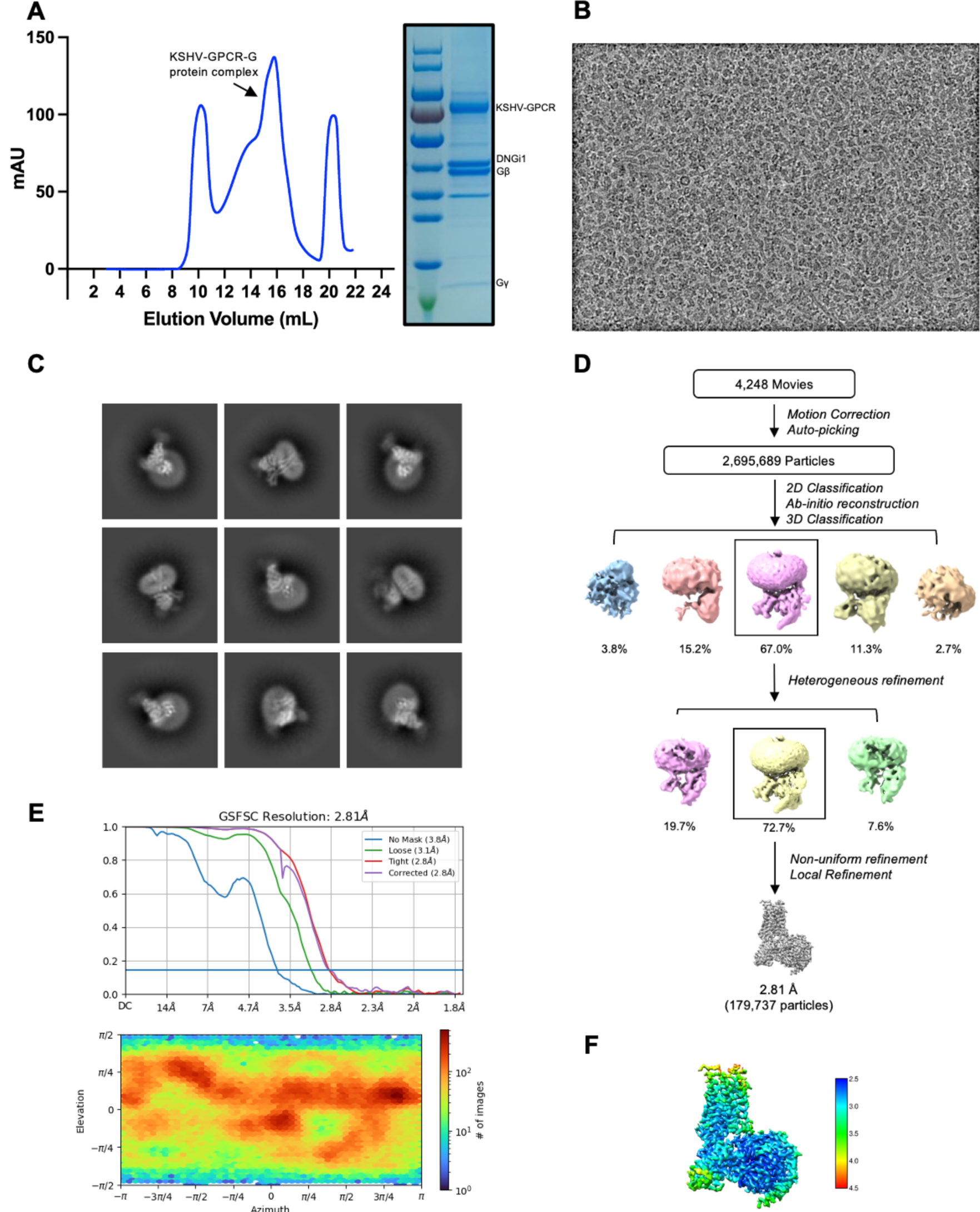
Protein purification, cryo-EM data collection, structure determination and cryo-EM maps of the KSHV-GPCR-G protein complex. **(A)** Size-exclusion chromatography elution profiles of the KSHV-GPCR-G protein complex and the SDS-PAGE and Coomassie blue staining of the KSHV-GPCR-G protein complex. **(B)** Representative micrograph of the complex particles from 4,248 movies. **(C)** Representative 2D averages. **(D)** Workflow for cryo-EM image processing. **(E)** Gold standard Fourier shell correlation (FSC) curve indicates overall nominal resolution at 2.81 Å using the FSC=0.143 criterion. **(F)** Local resolution map.

**Fig. S2.**
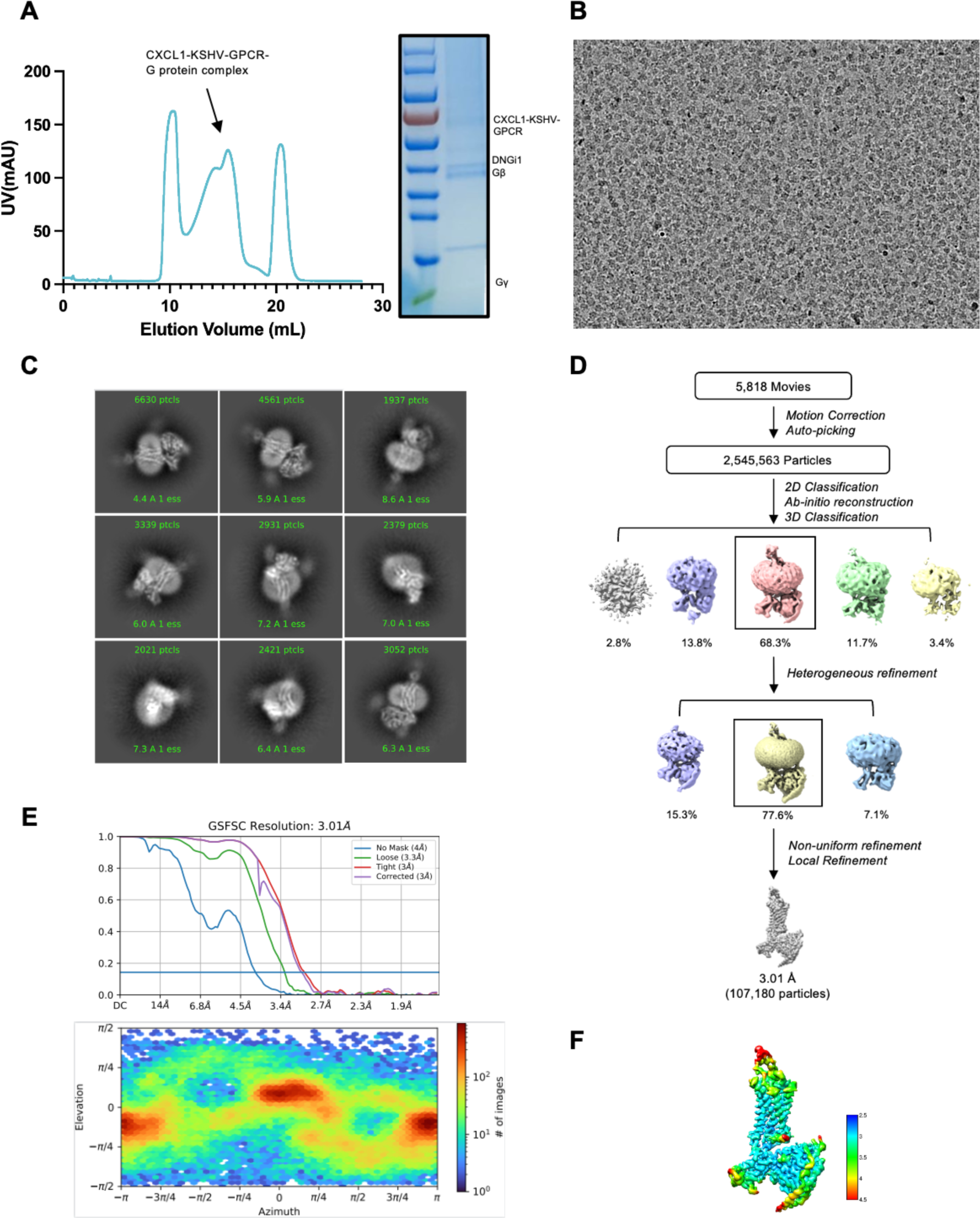
Protein purification, cryo-EM data collection, structure determination and cryo-EM maps of the CXCL1-KSHV-GPCR-G protein complex. **(A)** Size-exclusion chromatography elution profiles of the CXCL1-KSHV-GPCR-G protein complex and the SDS-PAGE and Coomassie blue staining of the CXCL1-KSHV-GPCR-G protein complex. **(B)** Representative micrograph of the complex particles from 5,818 movies. **(C)** Representative 2D averages. **(D)** Workflow for cryo-EM image processing. **(E)** Gold standard Fourier shell correlation (FSC) curve indicates overall nominal resolution at 3.01 Å using the FSC=0.143 criterion. **(F)** Local resolution map.

**Fig. S3.**
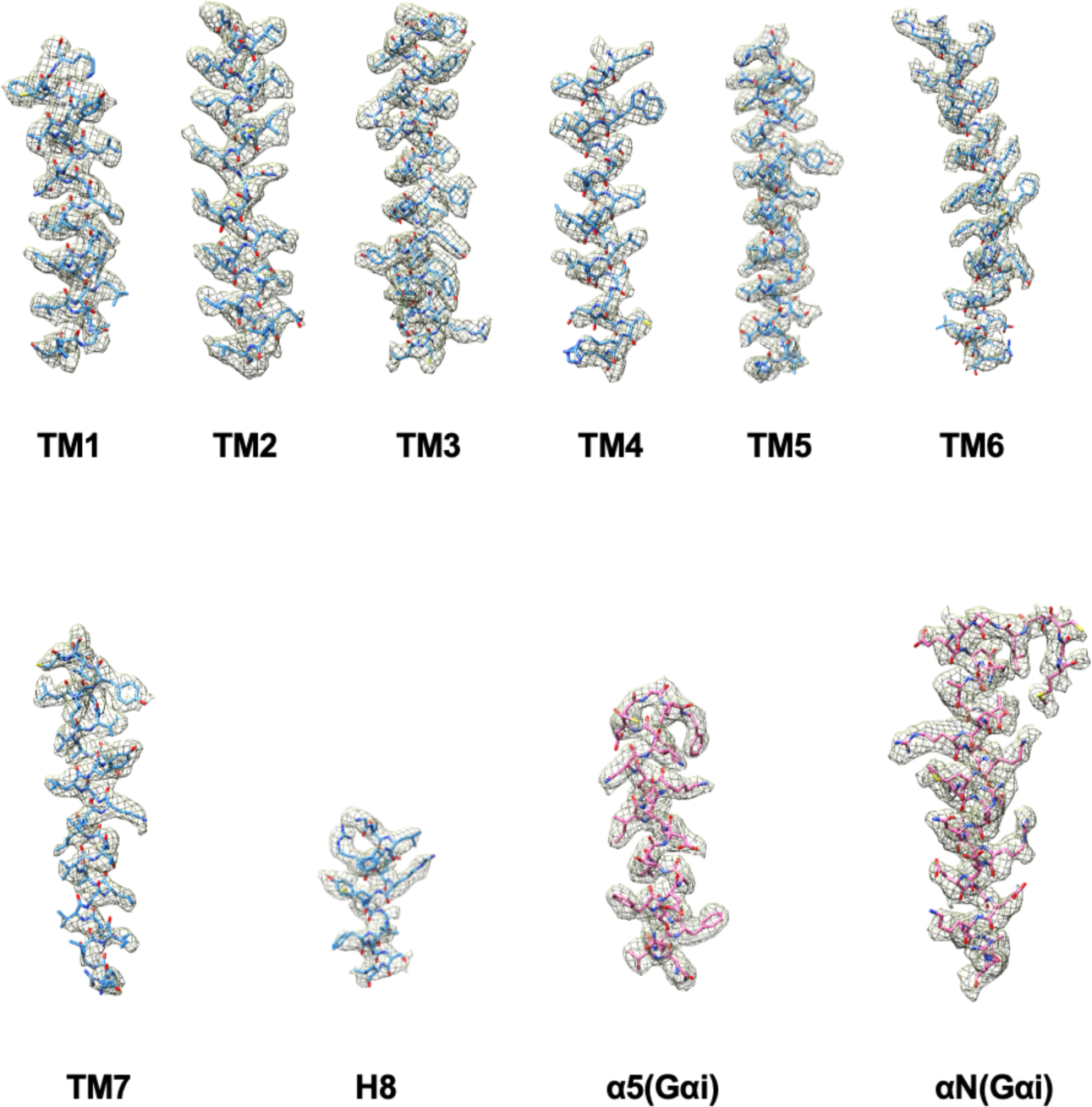
Representative density maps and models for TM1-7 and H8 of KSHV-GPCR and C-terminal α helices of Gαi1 (α5 and αN).

**Figure S4.**
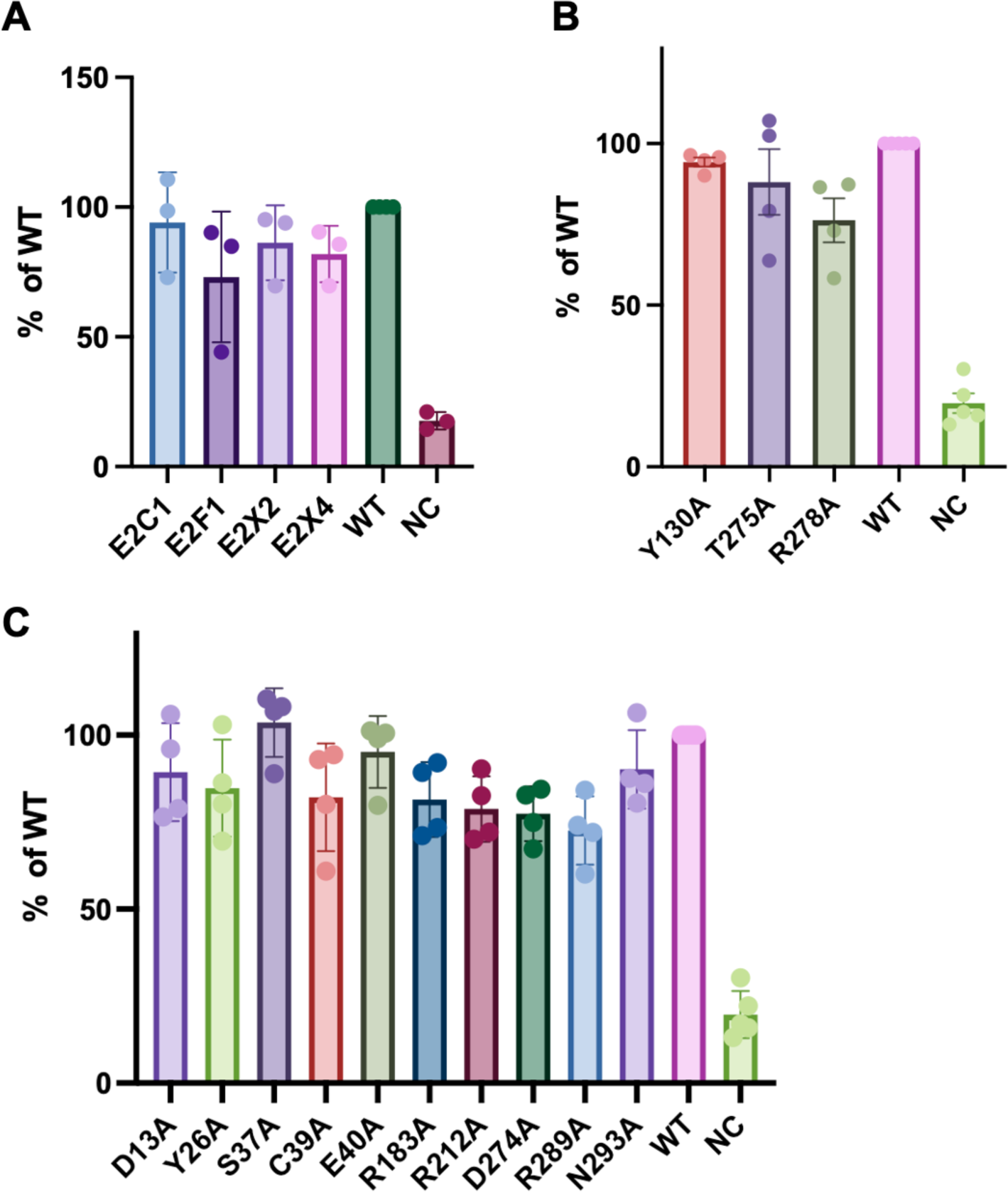
Cell surface expression of mutants. FLAG-tagged WT or mutant KSHV-GPCR (**A** ECL2 substitution, **B** ECL2 interacting residues and **C** TM binding pocket) were transiently expressed in HEK293T cells for 24 h. Cells were then incubated with an FITC-conjugated FLAG antibody for 30 min on ice. The mean fluorescence intensity signals were detected by flow cytometry. Data shown are means ± SEM from 3 independent experiments. *, *p* < 0.05.

## Notes

### Competing Interest Statement

The authors have declared no competing interest.

